# Genomic characterisation of *Listeria monocytogenes* isolated from normally sterile human body fluids in Lithuania from 2016 to 2021

**DOI:** 10.1101/2025.01.23.633655

**Authors:** Anželika Slavinska, Elita Jauneikaite, Ugnė Meškytė, Agnė Kirkliauskienė, Adam Misevič, Aurelija Petrutienė, Nomeda Kuisiene

## Abstract

*Listeria monocytogenes* is a saprophytic gram positive bacterium and opportunistic foodborne pathogen that can cause listeriosis in humans. The incidence of listeriosis has been rising globally and, despite antimicrobial treatment, the mortality rates associated with the most severe forms of listeriosis such as sepsis, meningitis, and meningoencephalitis remain high. The notification of listeriosis in humans is mandatory in Lithuania and up to 10 cases are reported annually. However, no studies have described the detailed virulence and antimicrobial susceptibility profiles of any clinical *L. monocytogenes* strains in Lithuania. Accordingly, this study aimed to describe the antibiotic susceptibility of invasive *L. monocytogenes* and perform in-depth characterisation of strains isolated from patients with neuroinfections through whole-genome sequencing.

A total of 70 isolates were collected, mostly from infected patients aged 65 or older, between 2016 and 2021: 41 (58.6%) from blood, 19 (27.1%) from cerebrospinal fluid, five (7.1%) from wounds, one (1.4%) from pleural fluid, and one (1.4%) from a brain abscess. Two phylogenetic lineages were identified—I (n = 16/70, 22.9%) and II (n = 54/70, 77.1%)—along with three serogroups—IIa (n = 53/70, 75.7%), IVb (n = 16/70, 22.9%), and IIc (n = 1/70, 1.4%). Genomic analysis of 20 isolates showed a high level of diversity with seven genotypes: ST6 (n = 6), ST155 (n = 5), ST8 (n = 4), ST504 (n = 2), and singletons for ST37, ST451, and ST2. Phylogenetic analysis clustered these isolates into two clades defined by serogroups IVb and IIa. Notably, five isolates were clustered tightly together (difference of 6–48 core SNPs from reference and 0, 4, or 44 SNPs from each other) with ST155, previously reported in a European outbreak. Comparison with publicly available *L. monocytogenes* genomes did not identify unique clusters or genotypes. No acquired antimicrobial resistance genes were identified.

Our study highlights the complementary value of whole-genome sequencing in routine PCR-based surveillance in Lithuania. This is the first study to characterise and compare genomes for *L. monocytogenes* associated with neuroinfections in Lithuania using whole-genome sequencing. The retrospective detection of the ST155 clone underscores the need for a review and strengthening of epidemiological surveillance strategies in clinical and non-clinical settings in Lithuania.

**Data summary:** The raw sequencing reads from the *L. monocytogenes* isolates generated in this study are available from the European Nucleotide Archive (https://www.ebi.ac.uk/ena/browser/home) under project accession number **PRJEB80562**; accession numbers for individual isolates are provided in **Suppl. Table S1**. Draft genomes of the *L. monocytogenes* strains were submitted to the BIGSdb-*Lm* database (https://bigsdb.pasteur.fr/listeria/) under accession numbers 110263–110281 and 108726.

## Introduction

*Listeria monocytogenes* is a foodborne pathogen that can cause various diseases, such as self-limiting gastroenteritis, sepsis, meningitis, encephalitis, and spontaneous abortion [1], with pregnant women, infants, immunocompromised individuals, and older adults demonstrating the worst outcomes [2]. Listeriosis is usually acquired via consumption of contaminated food [3] but is also widely present in nature in a range of animals, plants, and environments [4]. Globally, 0.1 to 10 cases per 1 million population of listeriosis are reported annually [5] with similar numbers reported in the EU (including the UK)—2.1 per 100,000 population [6]. Although the numbers of listeriosis cases are low, the infections have high rates of adverse outcomes and mortality [7,8], and its potential for causing foodborne outbreaks [9–11], makes *L. monocytogenes* an important public health threat.

The β-lactam antibiotics penicillin and aminopenicillins, including ampicillin and amoxicillin, are the first-line treatments for listeriosis [12,13]. β-lactam antibiotics are usually combined with an aminoglycoside, most commonly gentamicin [12]. Trimethoprim/sulfamethoxazole is used in cases of suspected penicillin allergy [14]. The use of fluoroquinolones, rifampicin, linezolid, and vancomycin has also been reported [15]. Unlike many other important human pathogens, *L. monocytogenes* has largely retained its susceptibility to antibiotics that have been used for decades in both humans and animals, such as penicillins [16]. *L. monocytogenes* is naturally resistant to specific antimicrobials, including third-generation cephalosporins and monobactams, because it does not have the appropriate penicillin-binding proteins [12] and is inherently resistant to first-generation quinolones, fosfomycin, cationic antimicrobial peptides, lincomycin, and sulfonamides due to intrinsic resistance genes, such as *norB* [17], *fosX* [18,19], *mprF* [20], *lin* [21], and *sul* [22]. The regulation of intrinsic resistance genes in *L. monocytogenes* is highly complex, making it difficult to predict phenotypes [23].

Various genomic and non-genomic methods have been applied to survey infections, identify potential outbreaks, and provide genotyping data on *L. monocytogenes* populations. *L. monocytogenes* has broadly been grouped into lineages I–IV [24]. Isolates from humans and food-sources predominantly fall into two phylogenetic lineages [25] consisting of serotypes 1/2a, 1/2b, and 4b, which are responsible for the majority of human listeriosis cases; serotype 4b has also been assigned to the leading hypervirulent clones of clonal complexes (CCs) 1, 2, 4, and 6 [2,26– 28]. Whole-genome sequencing (WGS) has become a widely used method for genotyping *L. monocytogenes* [29] and has proven useful in outbreak source investigations [30,31]; however, this method is rarely adopted for monitoring in countries with limited funding for epidemiological surveillance, such as Lithuania, where the notification of listeriosis in humans is mandatory and up to 10 cases are reported annually (0.21-0.46 per 100,000 populations between 2019-2022) [6].

Notably, there have been no detailed studies on the virulence and antimicrobial susceptibility profiles of clinical *L. monocytogenes* strains circulating in Lithuania. Hence, this study assessed the antimicrobial susceptibility of clinical *L. monocytogenes* strains in Lithuania and characterised in detail isolates from neuroinfection cases using WGS.

## Materials and Methods

### Bacterial isolates

We examined 70 *L. monocytogenes* isolates provided by the National Public Health Surveillance Laboratory (Vilnius, Lithuania). The isolates were collected from different clinical samples between 10 October 2016 and 20 September 2021 (**Suppl. Table S1**). Pure *L. monocytogenes* cultures were stored at -80 °C in brain heart infusion broth (Liofilchem, Roseto degli Abruzzi, Italy) containing 20% glycerol. To recover the isolates from long-term storage, 10 μL of stock was cultured on sheep blood agar plates (bioMérieux, Marcy-l’Étoile, France) at 37 °C for 24–48 h. A single colony from a visually pure culture of each *L. monocytogenes* isolate was selected for analysis.

### Antibiotic susceptibility testing

Minimum inhibitory concentrations (MICs) for penicillin G (0.063–4.0 mg/L), ampicillin (0.25–16.0 mg/L), erythromycin (0.125–4.0 mg/L), meropenem (0.25–16.0 mg/L), tetracycline (0.5–4.0 mg/L), and vancomycin (1.0–8.0 mg/L) were determined using the MICRONAUT-S (Sifin diagnostics Gmbh, Berlin, Germany) broth microdilution method and interpreted according to the European Committee on Antimicrobial Susceptibility Testing (EUCAST) breakpoints v14.0 [32]. The EUCAST provides MIC breakpoint values for *L. monocytogenes* only for benzylpenicillin, ampicillin, meropenem, and erythromycin. For other antibiotics, MICs were interpreted according to the EUCAST breakpoints for *Staphylococcus spp*. as previously described [33,34]. Antimicrobial susceptibility testing was performed using MICRONAUT-H Medium for fastidious bacteria (Sifin diagnostics Gmbh) following incubation at 35–37 °C in 5% CO_2_ for 22–24 h. For quality control, *Streptococcus pneumoniae* ATCC 49619 was used.

### Genomic DNA extraction

Genomic DNA was extracted from single bacterial colonies using a GeneJET Genomic DNA Purification Kit (Thermo Fisher Scientific, Waltham, MA, USA), following the manufacturer’s protocol. DNA quality was assessed by measuring absorbance at 260/280 nm with a BioPhotometer (Eppendorf, Hamburg, Germany), and DNA integrity was verified using 1% agarose gel electrophoresis.

### PCR characterisation of isolates

Genes specific for the genus *Listeria* (*prs*), species *monocytogenes* (*isp*), lineages LI and LII (*L1, L2*), and serotypes (ORF2819, ORF2110, *lmo0737, lmo1118*) of *L. monocytogenes* were targeted using PCR to confirm the presence of *L. monocytogenes* and define the specific lineages as previously described [35,36]. Amplification products were visualised on 1% agarose gel.

### Whole-genome sequencing and bioinformatics analyses

Genomic DNA for 20 *L. monocytogenes* isolates from cerebrospinal fluid samples was sent to MicrobesNG for whole-genome sequencing (https://microbesng.com/). DNA libraries were prepared using a Nextera XT DNA Library Preparation Kit (Illumina, San Diego, CA, USA) following the manufacturer’s protocol and sequenced on a NovaSeq 6000 system (Illumina) using a 250-bp paired-end protocol.

*Quality Control*. The quality of the raw reads was assessed using FastQC v0.12.1 (https://github.com/s-andrews/FastQC) and MultiQC v1.21 (https://github.com/MultiQC/MultiQC). Trimmomatic v0.39 (https://github.com/usadellab/Trimmomatic) [37] was used to trim low quality bases using the following parameters: LEADING: 20, TRAILING: 20, SLIDINGWINDOW: 5:30, MINLEN: 50. Bacterial species was confirmed using Kraken2 v2.1.3 (https://github.com/DerrickWood/kraken2) [38] and Bracken v2.9 (https://github.com/jenniferlu717/Bracken) [39] with the Kraken database (last updated 11/11/2023). Reads were *de novo* assembled using Spades v3.15.5 (https://github.com/ablab/spades) [40]. Draft genomes were assessed using Quast v5.2.0 (https://github.com/ablab/quast) [41] and CheckM2 v1.0.2 (https://github.com/chklovski/CheckM2) [42]. Bioawk v1.0 (https://github.com/lh3/bioawk) was used to remove contigs of ≤200 bp. A summary of the assembly statistics is provided in **Suppl. Table S2**.

*Characterisation*. Abricate v1.0.0 (https://github.com/tseemann/abricate) was used to detect acquired antimicrobial resistance genes using the Resfinder database (last updated 18/01/2023) [43]. The *Listeria monocytogenes* MLST database hosted by the Institut Pasteur (BIGSdb-*Lm*) (https://bigsdb.pasteur.fr/listeria/) [44] was used to detect intrinsic antimicrobial resistance genes. Genotypes were assigned using mlst v2.23.0 (https://github.com/tseemann/mlst) and BIGSdb-*Lm* [44]. MOB-suite v3.1.8 (ttps://github.com/phac-nml/mob-suite) [45] was used to reconstruct and characterise plasmids from draft assemblies. Virulence factor profiles were determined using the Virulence Factor Database [46] and BIGSdb-*Lm* [44]. PHAge Search Tool Enhanced Release (PHASTEST) [47] was used to assess the presence of prophage sequences. Incomplete prophages were excluded from analysis. The coding sequences and non-coding RNAs were predicted using Bakta v1.9.4 (https://github.com/oschwengers/bakta) [48].

*Phylogenetic analysis*. Snippy v4.6.0 (https://github.com/tseemann/snippy) was used to call SNPs from the whole genome sequences of 20 *L. monocytogenes* isolates using the recent ST155 genome (accession number: ERR7113321 [49], PubMLST assembly ID: 103485_20_06303) as reference. Recombination analysis was done using Gubbins v3.3.5 [50]. Recombination-free core SNPs were used to build a phylogenetic tree using IQ-TREE v2.0.3 [51] with the ‘GTR+G+ASC’ model, which was visualised and annotated using iTOL v6.9.1 (https://itol.embl.de/about.cgi) [52].

*Comparative analysis*. We compared our WGS results to 317 assembled genomes, downloaded from BIGSdb-*Lm* (https://bigsdb.pasteur.fr/listeria/), of isolates collected internationally (**Suppl. Table S3**). A phylogenetic tree was constructed as described in the phylogenetic analysis. We compared the virulence profiles of *L. monocytogenes* isolates from Lithuania with those for a subset of 22 assembled genomes. The subset consisted only of genomes that had ≤30 allelic differences (and <300 contigs) with Lithuanian strains selected from a minimum spanning tree (MST) created using the GrapeTree tool and BIGSdb-*Lm* (https://bigsdb.pasteur.fr/listeria/) (**Suppl. Fig. S1**). The GrapeTree was constructed based on the cgMLST1748 scheme and was color-coded according to the country of isolation [53].

## Results

### Overview of clinical *L. monocytogenes* strains: typing and antimicrobial susceptibility

A total of 70 *L. monocytogenes* isolates collected between 2016 and 2021 were analysed in this study. Among them, 41 (58.6%) were isolated from blood, 19 (27.1%) from cerebrospinal fluid, 5 (7.1%) from wounds, 1 (1.4%) from pleural fluid, and 1 (1.4%) from a brain abscess (**Suppl. Table S1**). The highest number of isolates was recorded in 2020 (n = 23, 32.9%). The majority of isolates were from patients aged 65 years and older (n = 35, 50%), followed by the 45–64 (n = 19, 27.1%) and 25–44 age groups (n = 6, 8.6%); there were only 3 (4.3%) isolates from patients younger than 1 year. Single cases were identified in the 1–4 and 15–24 age groups (1.4% each). The ages of five patients were unknown.

All isolates (n = 70) expressed the *prs* and *isp* genes, confirming their identity as *L. monocytogenes*. We identified two phylogenetic lineages—I (n = 16/70, 22.9%) and II (n = 54/70, 77.1%)—and three serogroups—IIa (n = 53/70, 75.7%), IVb (n = 16/70, 22.9%), and IIc (n = 1/70, 1.4%) (**Suppl. Table S1**). Serogroup IIa was found in isolates for 2016–2021, including all eight isolates in the 2021 collection (**Suppl. Table S1, Suppl. Fig. S2A**).

There was no significant difference in serogroup distributions between different age groups. Serogroup IIa was present in all age groups and was the only serogroup in patients aged less than 1 to 24 years. The highest serogroup diversity was observed in the 45–64 age group (n = 19), where serogroup IIa accounted for 11 cases (57.9%), serogroup IVb for seven cases (36.8%), and serogroup IIc for one case (5.3%). The largest number of cases was reported in the ≥65 years group (n = 35), with 29 cases (82.9%) associated with serogroup IIa and six (17.1%) with serogroup IVb (**Suppl. Fig. S2B**).

All *L. monocytogenes* isolates were susceptible to penicillin G, ampicillin, and erythromycin. Only four isolates (5.7%) were susceptible to tetracycline (MIC = 1) and the rest had MICs of 2 to ≥4 mg/L and would be deemed resistant. Based on the breakpoints for *S. aureus* (S ≤ 2 mg/L; R > 2 mg/L), three isolates (4.3%) were found to be resistant to vancomycin (**Suppl. Table S1**).

### Genomic characterisation of 20 *L. monocytogenes* isolates from neuroinfections (cerebrospinal fluid and brain abscess samples)

We investigated the genomic diversity of *L. monocytogenes* isolated from the cerebrospinal fluid (n = 19) and a brain abscess (n = 1) of patients with neuroinfections. The characteristics of the assembled genomes are listed in **Suppl. Table S2**.

Our analysis revealed seven distinct sequence types (STs), seven cgMLST sublineages (SLs), and 16 distinct cgMLST types (CTs) (**Suppl. Table S2**). The most common ST was ST6 (n = 6), followed by ST155 (n = 5), ST8 (n = 4), and ST504 (n = 2), while ST37, ST451, and ST2 were found in one isolate each (**Suppl. Table S2**). Phylogenetic analysis (**Fig. 1**) showed two clades that were separated based on the PCR serogroup: serogroup IVb and IIa isolates were grouped separately. Notably, five isolates were clustered with ST155 (PubMLST assembly ID: 103485_20_06303), which was previously reported in an outbreak in Europe [49].

**Figure 1.**
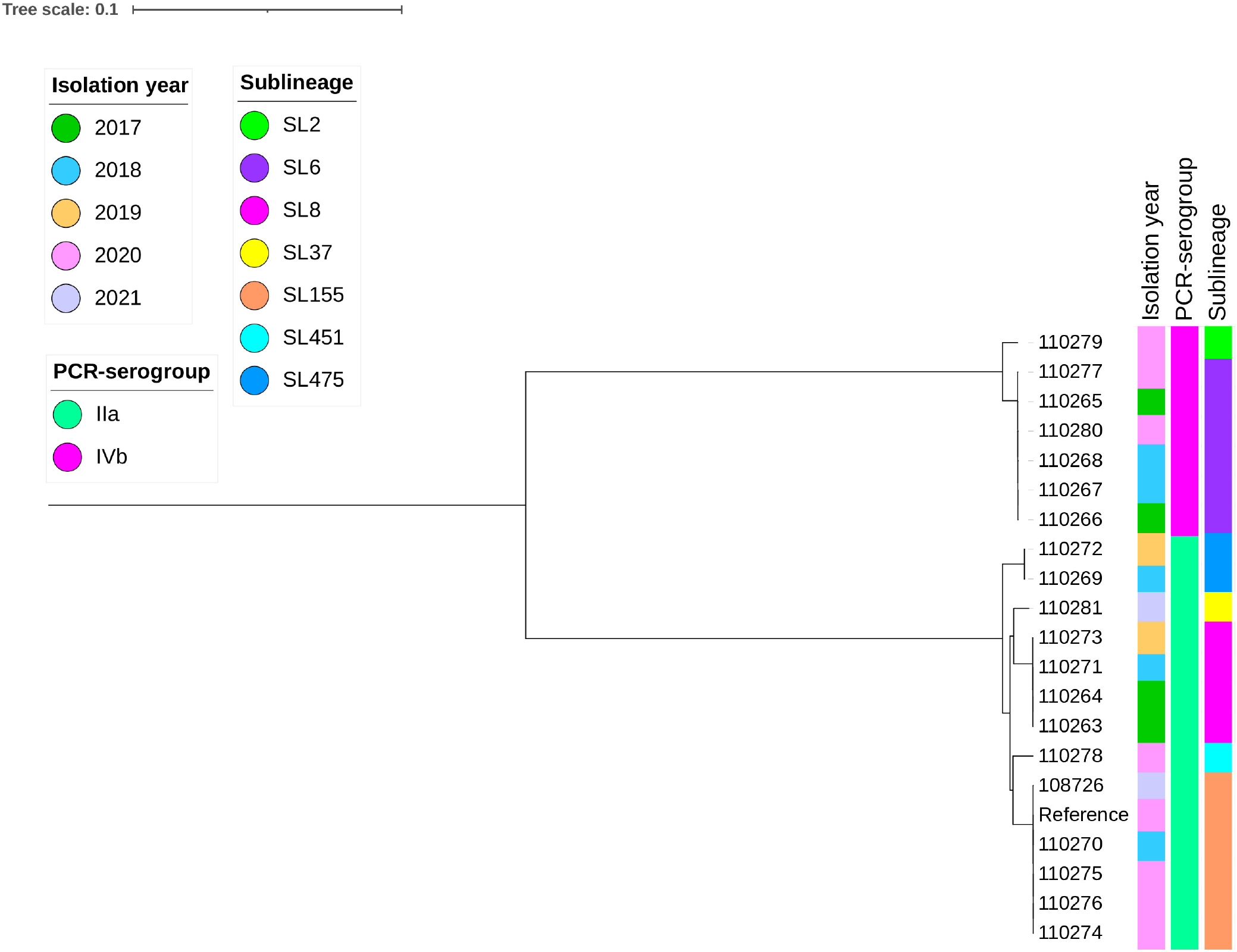
Phylogenetic relationship between 20 *L. monocytogenes* isolates from Lithuania. Maximum likelihood phylogenetic tree based on core SNPs of *L. monocytogenes* using the recent ST155 genome (PubMLST assembly ID: 103485_20_06303). Two clades were detected, consistent with the PCR serogroups and sublineages. Five isolates were clustered (6–48 core SNP difference) with the reference genome, isolated during a recent European outbreak [49]. Tree scale indicates nucleotide substitution per site.

The isolates differed only by 6–48 core SNPs from the reference and 0, 4, or 44 SNPs from each other, with isolate 110270 from 2018 differing by six SNPs from the reference and only 4 SNPs from isolates 110274, 10276, and 110275. The latter isolates were collected within 2–3 months of each other and differed only by six SNPs different from the reference and 0 SNPs between themselves, suggesting that the ST155 *L. monocytogenes* clone has spread beyond the initial outbreak (the other closest isolate to this clade 110278 was 3,665 SNPs different from the reference).

Among the relevant virulence genes, stress survival islet 1 (SSI-1), associated with tolerance to acidic, bile, gastric, and salt stresses, was detected in 45% (n = 9/20) of the isolates, all from lineage II. Seven isolates from lineage I and two from lineage II, harboured only *lmo0447* from SSI-1. SSI-2, associated with survival under high pH and oxidative stresses, was detected in two Lithuanian isolates from lineage II (110269 and 110272). All 20 isolates harboured the *lmo1800* gene, which encodes the phosphatase LipA, essential for promoting infections *in vivo* [54], and *lmo1799*, the putative peptidoglycan-binding protein gene (with a LPXTG motif) (**Fig. 2**). LPXTG surface proteins play crucial roles in the adhesion and invasion of *L. monocytogenes* [55]. The other annotated genes were functionally categorized into subsystems (**Suppl. Table S4**). No genes associated with phosphorus metabolism were detected in any strain.

**Figure 2.**
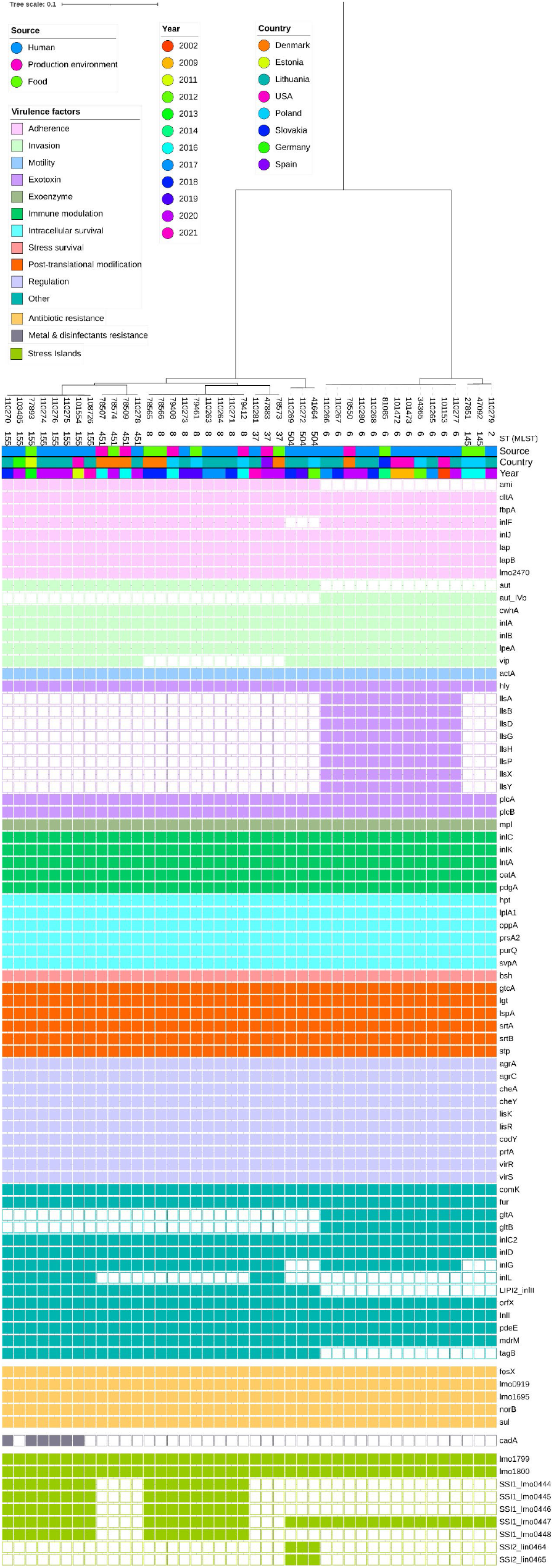
Distribution of virulence, antimicrobial resistance, and heavy metal resistance genes among *L. monocytogenes* isolates from Lithuania and genetically related strains from other countries. Maximum likelihood phylogenetic tree of 42 closely related genomes, including isolates clustered closely with those from Lithuania, based on core SNPs of *L. monocytogenes* using the recent ST155 genome (PubMLST assembly ID: 103485_20_06303). Tree scale indicates nucleotide substitution per site.

No acquired antimicrobial resistance genes were detected in any of the 20 genomes; only *fosX* (lmo1702), *lin* (lmo0919), *mprF* (lmo1695), *norB* (lmo2818), and *sul* (lmo0224) were identified, which mediate intrinsic resistance (**Suppl. Table S2, Fig. 2**).

### Mobile genetic elements and prophages in *L. monocytogenes*

We detected key plasmids in the assemblies of two isolates, 110263 and 110273, both belonging to ST8. The plasmid in 110263 showed the closest similarity to the plasmid with identifier LR134399 from *L. monocytogenes* strain NCTC7974; this plasmid was 291,717 bp in size, had 300 predicted coding sequences, and a GC content of 37.8% but was identified as a non-mobilizable plasmid through a MOBsuite analysis. The plasmid had genes related to virulence and stress responses (*adhR, iap, mogR, ltaP, secA, cidC, hdeD*, and *hflX*), motility and chemotaxis (*fliG, fliF, fliE, fliM, fliJ, fliN, flgC, flgB, flgE, cheA, cheV, motA*, and *motB)*, amino acid metabolism and transport, and noncoding RNAs (*rli31, rli32, rli33*, and *rli34*) (**Suppl. Table S5**).

Isolate 110273 had a plasmid very similar to pLMR479a [56], and was 86,759 bp long, with a GC content of 37.0%, and encoded a Group 2 RepA. The plasmid contained 85 predicted coding sequences (**Suppl. Table S6**). The plasmid expressed *cadA1, cadA2*, and *cadC*, which are associated with resistance to cadmium and other heavy metals [57], as well as *copB* and *copZ*, involved in copper homeostasis and resistance [58–60]. The plasmid also contained the virulence and adaptation genes *virB4, traM, fetA, fetB*, and *fixK* and the stress response genes *dps* and *dinB* [61,62].

We identified 11 distinct intact prophages. The most common prophages were *Listeria phage A118* (n = 14) and *Listeria phage vB_LmoS_188* (n = 11); the rarest were *Listeria phage B054* (n = 1) and *Listeria phage LP-030-2* (n = 1) (**Suppl. Table S7**). All ST155 Lithuanian genomes had an intact *Listeria phage LP-HM00113468*.

### Comparative genomics of Lithuanian and international *L. monocytogenes*

To put our *L. monocytogenes* genomes into a broader geographical context, we compared the Lithuanian *L. monocytogenes* genomes with those from a publicly accessible dataset of *L. monocytogenes* genomes provided by the Institut Pasteur. Using the GrapeTree MST and cgMLST1748 scheme, we identified 297 closely related isolates (317 including the Lithuanian isolates) (**Suppl. Fig. S1**).

These isolates came from various sources: 150 from clinical samples (47.3%), 115 from food products (36.3%), 42 from food processing environments (13.2%), three from animals (0.9%), one from the natural environment (0.3%), and six with unknown origins, confirming the range of environments *L. monocytogenes* can adapt to. The dataset primarily comprised isolates from Poland (n = 126, 39.7%), Denmark (n = 41, 12.9%), the USA (n = 39, 12.3%), and Lithuania (n = 20, 6.3%), with ≤10 isolates collected from other countries (**Suppl. Fig. S1**). SNP-based phylogenetic analysis revealed a distinct cluster of isolates that formed clonal complex (CC) 155 (**Fig. 3**). CC155 contains isolates from humans, as well as food and food production environments; this provides evidence that food-to-human transmission is occurring in these *L. monocytogenes* infections (**Fig. 3**). The other *L. monocytogenes* isolates were clustered near other isolates from human and food sources (**Fig. 3**). Overall, *L. monocytogenes* genomes in Lithuania were clustered with genomes from various locations and years.

**Figure 3.**
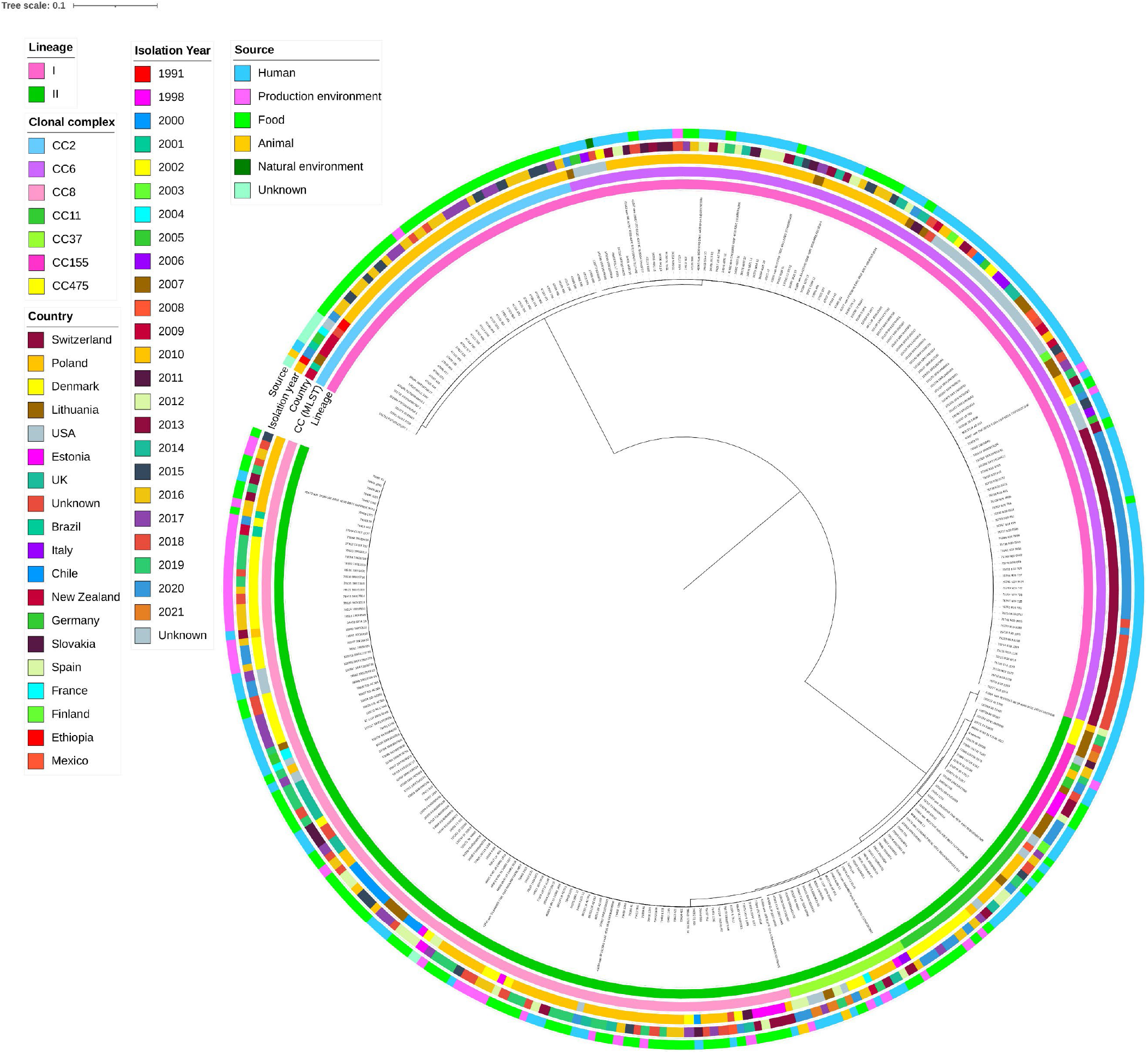
Comparative phylogenetic tree of Lithuanian and other *L. monocytogenes* genomes available from the Institut Pasteur. Maximum likelihood phylogenetic tree of 314 *L. monocytogenes* genomes, including 20 genomes from this study, based on core SNPs of *L. monocytogenes* using the recent ST155 genome (PubMLST assembly ID: 103485_20_06303). Tree scale indicates nucleotide substitution per site.

We compared the distribution of virulence and antimicrobial resistance genes between the Lithuanian and 22 closely related *L. monocytogenes* genomes (**Fig. 2**), based on the cgMLST and phylogenetic analyses. Of the 93 virulence genes, 72 (77.4%) distinct genes were detected among isolates (min = 58, max = 67). No significant differences in virulence gene profiles were observed between the isolates from different countries or years, confirming that specific virulence gene profiles are more likely to be associated with *L. monocytogenes* clones than geographical distribution. In total, 30/42 (71.4%) isolates had the *vip* gene, with 22/30 (73.3%) coming from humans (**Fig. 2**). A total of 12/42 isolates (28.6%) carried the LIPI-3 pathogenicity island, which was most frequently detected in isolates from humans (n = 10/27). The adhesion-associated gene *ami*, invasion gene *aut*, LIPI-2 pathogenicity island-associated gene *inlII*, and teichoic acid biosynthesis-related gene *tagB* were found in all serogroup IIa strains but were absent in serogroup IVb strains. Meanwhile, *aut-IVb, gltA*, and *gltB* and genes within the LIPI-3 pathogenicity island (*llsA, llsB, llsD, llsG, llsH, llsP, llsX*, and *llsY*) were only found in serogroup IVb strains, with *aut-IVb, gltA*, and *gltB* present in all serogroup IVb strains (n = 15). All strains carried the LIPI-1 pathogenicity island, which is critical for *L. monocytogenes* infection. Overall, five virulence gene alleles were identified only in Lithuanian strains (**Table 1**).

**Table 1.**
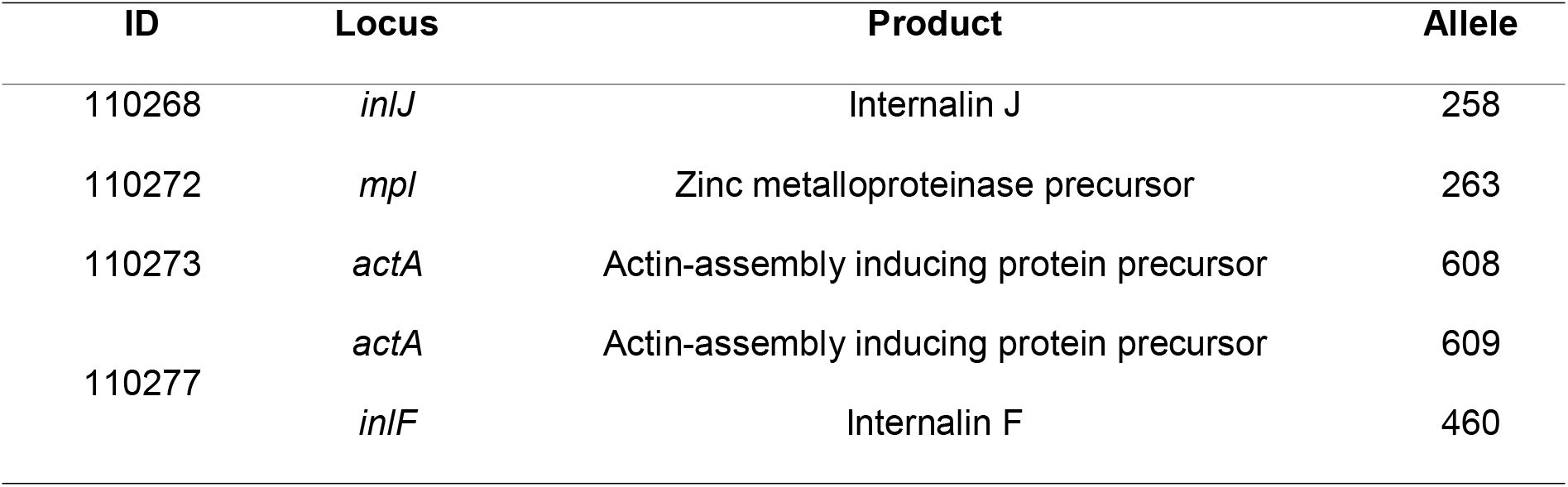
Unique alleles of virulence genes found only in Lithuanian *L. monocytogenes* strains.

| ID | Locus | Product | Allele |
| --- | --- | --- | --- |
| 110268 | *inlJ* | Internalin J | 258 |
| 110272 | *mpl* | Zinc metalloproteinase precursor | 263 |
| 110273 | *actA* | Actin-assembly inducing protein precursor | 608 |
| 110277 | *actA* | Actin-assembly inducing protein precursor | 609 |
|  | *inlF* | Internalin F | 460 |

We only identified one gene associated with resistance to heavy metals and disinfectants. The gene *cadA*, which encodes a P-type ATPase and is associated with resistance to heavy metals (cadmium) [63], was found exclusively in six isolates belonging to SL155, including four isolates from Lithuania (110270, 110276, 110275, and 110274), one from the USA (101554), and one from Estonia (77893). We analysed the homologous sequences of the *cadA* gene among the Lithuanian *L. monocytogenes* and other bacteria using BLASTn and identified highly conserved (>99.9%) homologous sequences in *Enterococcus* spp. (e.g. *E. faecium, E. faecalis*, and *E. saigonensis)*.

## Discussion

In recent years, WGS has become the primary tool for epidemiological surveillance in national programs, outbreak investigations, and environmental monitoring in food processing facilities, greatly enhancing food safety controls and public health protection [64–66]. Genomic analysis allows for the rapid classification of bacterial isolates and provides information on the potential phylogenetic relationships. Genomic epidemiology is useful for tracking not only local outbreaks but also those involving multiple institutions and countries [49]. However, in Lithuania, high-resolution methods with strong discriminatory power are not yet implemented in routine listeriosis surveillance. Instead, routine monitoring primarily relies on methods with lower discriminatory capacity, such as serotyping using agglutination reactions or serogroup prediction using PCR-based techniques. In this study, we performed a detailed characterisation of 20 *L. monocytogenes* isolates from neuroinfection cases using WGS and assessed the antimicrobial susceptibility of clinical isolates collected between 2016 and 2021.

In the EU/EEA countries, including the UK, over 2,300 cases of listeriosis are reported annually (the lowest number was observed in 2020, with 1,928 cases) [67]. The reported cases in 2022 were the highest in over 10 years, with 2,772 cases. The age group most affected was those over 64 years, with 1,903 cases (71.0%) and a notification rate of 2.1 per 100,000 population [6]. Notably, Lithuania exhibited a high case fatality rate (average: 45.9%, peaking in 2017 at 80%), contrasting sharply with the that in the EU/EEA countries (average: 15.9%, reaching a minimum of 14.1% in 2017) [67]. However, little is known about the population structure of *L. monocytogenes* from sporadic cases in Lithuania. This study showed that more cases of listeriosis were reported in 2020 and, overall, more cases were detected in patients aged 65 years old and older.

Molecular characterisation of *L. monocytogenes* isolates (n = 70) from clinical samples in Lithuania identified three serogroups: serogroup IIa and IIc, lineage II (n = 54/70); and serogroup IVb, lineage I (n = 16/70). This aligns with the findings of previous reports indicating that the majority of human cases are caused by lineages I (serotypes 1/2b and 4b) and II (serotype 1/2a) [15].

We evaluated the antimicrobial susceptibility of 70 *L. monocytogenes* strains using a phenotypic assay based on the standardized broth dilution method, which is considered the gold standard for detecting antimicrobial resistance [68]. All isolates were susceptible to clinically relevant antibiotics, including penicillin G, ampicillin, and erythromycin.

WGS analysis of 20 isolates from Lithuania showed that the most common ST was ST6 (n = 6), followed by ST155 (n = 5), ST8 (n = 4), ST504 (n = 2), while ST37, ST451, and ST2 were found in one isolate each. Among these, ST6, ST2, ST8, ST37, and ST451 have been associated with invasive listeriosis presenting as meningitis and septicaemia [69–73]. In contrast, ST8, ST155, and ST504, while implicated in *L. monocytogenes* infection, have predominantly been isolated from food products, food processing environments, and processing water [74,75].

Genomic comparison of the Lithuanian *L. monocytogenes* genomes with similar publicly available genomes showed that the Lithuanian genomes were intermixed with those from other countries, mainly European countries (Poland, Estonia, Denmark, Slovakia, Germany, and the USA), isolated from humans, food products, or food-processing environments. This is not surprising given the strong relationships reported by an increasing number of molecular epidemiology studies between different sources and geographical areas [49]. Notably, the five Lithuanian ST155 genomes identified in this study were closely related to a strain isolated in Germany that belongs to the Omikron1 cluster (ENA code: ERR7113321; PubMLST ID: 103485) and has been associated with an outbreak in Europe [49]. This Omikron1 strain was classified as serogroup IIa, CC155/SL155, and CT842/CT5098, and the outbreak originated from two food-processing facilities in Lithuania [49]. Omikron1 subcluster 1 strains have been linked to 64 confirmed listeriosis cases from 2016 to 2023 in Austria, Belgium, Germany, Italy, and the Netherlands, with 10 fatalities, predominantly occurring in 2020 [49]. Unfortunately, we did not have clinical or detailed metadata on the patients infected with ST155 in this study and could thus not make further extrapolations on these cases and the reported outbreak in Europe.

Notably, we identified an isolate with a plasmid expressing *cadA1, cadA2*, and *cadC*, which are associated with resistance to cadmium and other heavy metals [57], as well as *copB* and *copZ*, involved in copper homeostasis and resistance [58–60].

*L. monocytogenes* often exhibits tolerance to heavy metals and biocides. The overuse of disinfectants and cadmium resistance may enhance *L. monocytogenes* persistence in food products and food-processing environments [76]. Between 10% and 80% of *L. monocytogenes* strains isolated from food and food processing environments show tolerance to biocides [64]. The persistence of *L. monocytogenes* can elevate the risk of product contamination, food recalls, and foodborne outbreaks [77]. The identification of the ST155 outbreak strain from the environment of the two Lithuanian processing plants in 2023 and in products since 2016 reflects the high persistence of *L. monocytogenes* [49] and suggests that the sources of contamination have not been identified and properly controlled. Notably, cadmium resistance and plasmid carriage in *L. monocytogenes* occur more commonly among strains repeatedly isolated from foods than among those encountered only sporadically [78]. We did not identify any virulence-associated phages, though *Listeria* genomes often harbour prophages that contribute to genome variability and may enhance their adaptability and survival in various environments [79]. Further investigations of phages in these isolates should consider a long-read WGS using Oxford Nanopore technology to overcome the issue of incomplete page profiling, which is expected in short-read sequencing data, such as those used in this study. We identified unique virulence gene alleles in four Lithuanian strains: *inlJ258, mpl263, actA608, actA609*, and *inlF460*. However, to assess the potential impact of these alleles on pathogenicity and virulence, additional information is required, including details on disease progression, clinical outcomes, and relevant clinical data. This information was beyond the scope of our study, limiting the clinical interpretation of our findings.

## Conclusion

The *L. monocytogenes* isolates associated with neuroinfections in Lithuania between 2016 and 2021 were genetically diverse, with five isolates closely related to the ST155 genotype, which recently caused a multi-country outbreak associated with food production [49]. This indicates that the ST155 genotype was present in the population before the reported outbreak. This retrospective detection underscores the need for a review and strengthening of epidemiological surveillance strategies in Lithuania. Although we did not detect high levels of antimicrobial resistance, the emergence of antimicrobial resistance to first-line treatments, such as aminopenicillins (including amoxicillin, penicillin, and ampicillin), have been reported in a few countries [80]. Acquired antimicrobial resistance in *L. monocytogenes* is uncommon and the standard treatment for listeriosis remains effective. Ongoing surveillance in both clinical and food isolates is essential to identify the emergence of new resistance patterns and evaluate long-term trends in *L. monocytogenes* clone distributions in clinical and non-clinical settings.

## Author contributions

Conceptualization: AS, EJ, NK. Data curation: AS. Formal analysis: AS, EJ. Investigation: AS, EJ, UM, AK, AM. Methodology: AS, EJ, NK. Project administration: NK. Resources: AS, EJ, AK, AP, NK. Supervision: NK. Visualization: AS, EJ. Writing – original draft and further drafts: AS, EJ, NK. Writing – review & editing: All authors.

## Ethical approval

This study used strains obtained from the National Public Health Surveillance Laboratory (Vilnius, Lithuania). The Lithuanian Bioethics Committee waived the need for ethical clearance because it did not involve human biological samples or patient-related information. This retrospective study exclusively used pure cultures, with each sample encrypted and identified using a unique code to ensure anonymity. Under Lithuanian law, the use of microbial cultures does not fall under the scope of biomedical research. It is important to note that the isolates were collected as part of national pathogen and epidemiological surveillance, conducted in accordance with Order No. V-1194 issued by the Ministry of Health of the Republic of Lithuania on 18 December 2013.

## Acknowledgements

We would like to thank Editage (www.editage.com) for English language editing.

EJ is affiliated with the National Institute for Health Research Health Protection Research Unit (NIHR HPRU) in Healthcare Associated Infections and Antimicrobial Resistance at the Imperial College London in partnership with UK Health Security Agency (UKHSA) in collaboration with Imperial Healthcare Partners, the University of Cambridge, and University of Warwick.

We also thank the Institut Pasteur teams for the curation and maintenance of BIGSdb-Pasteur databases at http://bigsdb.pasteur.fr.

## References

[1] Magalhães R, Mena C, Ferreira V, Silva J, Almeida G, Gibbs P, et al. Listeria monocytogenes. Encycl. Food Safety, Second Ed. Vol. 1-4, vol. 1–4, StatPearls Publishing; 2023, p. V2-164-V2-178. 10.1016/B978-0-12-822521-9.00045-9.

[2] Maury MM, Tsai YH, Charlier C, Touchon M, Chenal-Francisque V, Leclercq A, et al. Uncovering Listeria monocytogenes hypervirulence by harnessing its biodiversity. Nat Genet 2016;48:308. 10.1038/NG.3501.

[3] Mughini-Gras L, Paganini JA, Guo R, Coipan CE, Friesema IHM, van Hoek AHAM, et al. Source attribution of Listeria monocytogenes in the Netherlands. Int J Food Microbiol 2025;427:110953. 10.1016/J.IJFOODMICRO.2024.110953.

[4] Lourenco A, Linke K, Wagner M, Stessl B. The Saprophytic Lifestyle of Listeria monocytogenes and Entry Into the Food-Processing Environment. Front Microbiol 2022;13:789801. 10.3389/FMICB.2022.789801/BIBTEX.

[5] Listeriosis n.d. https://www.who.int/news-room/fact-sheets/detail/listeriosis (accessed 14 December 2024).

[6] European Center for Disease Prevention and Control. Listeriosis, Annual Epidemiological report for 2022. Surveill Rep 2024:9.

[7] Charlier C, Perrodeau É, Leclercq A, Cazenave B, Pilmis B, Henry B, et al. Clinical features and prognostic factors of listeriosis: the MONALISA national prospective cohort study. Lancet Infect Dis 2017;17:510–9. 10.1016/S1473-3099(16)30521-7.

[8] Boelaert F, Stoicescu A, Amore G, Messens W, Hempen M, Rizzi V, et al. The European Union One Health 2019 Zoonoses Report. EFSA J 2021;19:e06406. 10.2903/J.EFSA.2021.6406.

[9] Halbedel S, Wilking H, Holzer A, Kleta S, Fischer MA, Lüth S, et al. Large Nationwide Outbreak of Invasive Listeriosis Associated with Blood Sausage, Germany, 2018-2019. Emerg Infect Dis 2020;26:1456–64. 10.3201/EID2607.200225.

[10] Fernández-Martínez NF, Ruiz-Montero R, Briones E, Baños E, Rodríguez-Alarcón LGSM, Chaves JA, et al. Listeriosis outbreak caused by contaminated stuffed pork, Andalusia, Spain, July to October 2019. Euro Surveill 2022;27. 10.2807/1560-7917.ES.2022.27.43.2200279.

[11] Stephan R, Horlbog JA, Nüesch-Inderbinen M, Dhima N. Outbreak of Listeriosis Likely Associated with Baker’s Yeast Products, Switzerland, 2022–2024. Emerg Infect Dis 2024;30:2424. 10.3201/EID3011.240764.

[12] Hof H, Nichterlein T, Kretschmar M. Management of listeriosis. Clin Microbiol Rev 1997;10:345–57. 10.1128/cmr.10.2.345.

[13] Grayo S, Join-Lambert O, Desroches MC, Le Monnier A. Comparison of the in vitro efficacies of moxifloxacin and amoxicillin against Listeria monocytogenes. Antimicrob Agents Chemother 2008;52:1697–702. 10.1128/AAC.01211-07.

[14] Koopmans MM, Brouwer MC, Vázquez-Boland JA, van de Beek D. Human Listeriosis. Clin Microbiol Rev 2023;36. 10.1128/CMR.00060-19.

[15] Baquero F F., Lanza V, Duval M, Coque TM. Ecogenetics of antibiotic resistance in Listeria monocytogenes. Mol Microbiol 2020;113:570–9. 10.1111/MMI.14454.

[16] Moura A, Leclercq A, Vales G, Tessaud-Rita N, Bracq-Dieye H, Thouvenot P, et al. Phenotypic and genotypic antimicrobial resistance of Listeria monocytogenes: an observational study in France. Lancet Reg Heal - Eur 2024;37:100800. 10.1016/j.lanepe.2023.100800.

[17] Ding Y, Onodera Y, Lee JC, Hooper DC. NorB, an efflux pump in Staphylococcus aureus strain MW2, contributes to bacterial fitness in abscesses. J Bacteriol 2008;190:7123–9. 10.1128/JB.00655-08/ASSET/75E3F838-C847-44C9-8DBC-A6978C734B34/ASSETS/GRAPHIC/ZJB0210881980004.JPEG.

[18] Fillgrove KL, Pakhomova S, Schaab MR, Newcomer ME, Armstrong RN. Structure and mechanism of the genomically encoded fosfomycin resistance protein, FosX, from Listeria monocytogenes. Biochemistry 2007;46:8110–20. 10.1021/BI700625P/SUPPL_FILE/BI700625P-FILE003.PDF.

[19] Scortti M, Han L, Alvarez S, Leclercq A, Moura A, Lecuit M, et al. Epistatic control of intrinsic resistance by virulence genes in Listeria. PLOS Genet 2018;14:e1007525. 10.1371/JOURNAL.PGEN.1007525.

[20] Thedieck K, Hain T, Mohamed W, Tindall BJ, Nimtz M, Chakraborty T, et al. The MprF protein is required for lysinylation of phospholipids in listerial membranes and confers resistance to cationic antimicrobial peptides (CAMPs) on Listeria monocytogenes. Mol Microbiol 2006;62:1325–39. 10.1111/J.1365-2958.2006.05452.X.

[21] Duval M, Dar D, Carvalho F, Rocha EPC, Sorek R, Cossart P. HflXr, a homolog of a ribosome-splitting factor, mediates antibiotic resistance. Proc Natl Acad Sci U S A 2018;115:13359–64. 10.1073/PNAS.1810555115/SUPPL_FILE/PNAS.1810555115.SD06.TXT.

[22] Moura A, Leclercq A, Vales G, Tessaud-Rita N, Bracq-Dieye H, Thouvenot P, et al. Phenotypic and genotypic antimicrobial resistance of Listeria monocytogenes: an observational study in France. Lancet Reg Heal - Eur 2023;37:100800. 10.1016/J.LANEPE.2023.100800.

[23] Krawczyk-Balska A, Markiewicz Z. The intrinsic cephalosporin resistome of Listeria monocytogenes in the context of stress response, gene regulation, pathogenesis and therapeutics. J Appl Microbiol 2016;120:251–65. 10.1111/JAM.12989.

[24] Orsi RH, Bakker HC de., Wiedmann M. Listeria monocytogenes lineages: Genomics, evolution, ecology, and phenotypic characteristics. Int J Med Microbiol 2011;301:79–96. 10.1016/J.IJMM.2010.05.002.

[25] Chenal-Francisque V, Maury MM, Lavina M, Touchon M, Leclercq A, Lecuit M, et al. Clonogrouping, a rapid multiplex PCR method for identification of major clones of listeria monocytogenes. J Clin Microbiol 2015;53:3355–8. 10.1128/JCM.00738-15.

[26] McLauchlin J. Distribution of serovars of Listeria monocytogenes isolated from different categories of patients with listeriosis. Eur J Clin Microbiol Infect Dis 1990;9:210–3. 10.1007/BF01963840.

[27] Farber JM, Peterkin PI. Listeria monocytogenes, a food-borne pathogen. Microbiol Rev 1991;55:476–511. 10.1128/mr.55.3.476-511.1991.

[28] Gray MJ, Zadoks RN, Fortes ED, Dogan B, Cai S, Chen Y, et al. Listeria monocytogenes isolates from foods and humans form distinct but overlapping populations. Appl Environ Microbiol 2004;70:5833–41. 10.1128/AEM.70.10.5833-5841.2004.

[29] Lakicevic B, Jankovic V, Pietzka A, Ruppitsch W. Wholegenome sequencing as the gold standard approach for control of Listeria monocytogenes in the food chain. J Food Prot 2023;86:100003. 10.1016/J.JFP.2022.10.002.

[30] World Health Organization. Whole genome sequencing for foodborne disease surveillance: landscape paper. 2018.

[31] World Health Organization. Whole genome sequencing as a tool to strengthen foodborne disease surveillance and response. Module 2. Whole genome sequencing in foodborne disease routine surveillance. Licence: CC BY-NC-SA 3.0 IGO. 2023. https://iris.who.int/handle/10665/373521 (accessed 11 December 2024).

[32] The European Committee on Antimicrobial Susceptibility Testing. Eur Comm Antimicrob Susceptibility Testing Break Tables Interpret MICs Zo Diameters Version 140 2024. https://www.eucast.org/clinical_breakpoints (accessed 6 August 2024).

[33] Noll M, Kleta S, Al Dahouk S. Antibiotic susceptibility of 259 Listeria monocytogenes strains isolated from food, food-processing plants and human samples in Germany. J Infect Public Health 2018;11:572–7. 10.1016/J.JIPH.2017.12.007.

[34] Prieto M, Martínez C, Aguerre L, Rocca MF, Cipolla L, Callejo R. Antibiotic susceptibility of Listeria monocytogenes in Argentina. Enferm Infecc Microbiol Clin 2016;34:91–5. 10.1016/J.EIMC.2015.03.007.

[35] Rawool DB, Doijad SP, Poharkar K V., Negi M, Kale SB, Malik SVS, et al. A multiplex PCR for detection of Listeria monocytogenes and its lineages. J Microbiol Methods 2016;130:144–7. 10.1016/J.MIMET.2016.09.015.

[36] Doumith M, Buchrieser C, Glaser P, Jacquet C, Martin P. Differentiation of the major listeria monocytogenes serovars by multiplex PCR. J Clin Microbiol 2004;42:3819–22. 10.1128/JCM.42.8.3819-3822.2004/ASSET/030268F4-5E8C-42B4-8931-5A6F6CE94BC0/ASSETS/GRAPHIC/ZJM0080444840001.JPEG.

[37] Bolger AM, Lohse M, Usadel B. Trimmomatic: a flexible trimmer for Illumina sequence data. Bioinformatics 2014;30:2114. 10.1093/BIOINFORMATICS/BTU170.

[38] Wood DE, Lu J, Langmead B. Improved metagenomic analysis with Kraken 2. Genome Biol 2019;20:257. 10.1186/S13059-019-1891-0.

[39] Lu J, Breitwieser FP, Thielen P, Salzberg SL. Bracken: Estimating species abundance in metagenomics data. PeerJ Comput Sci 2017;2017:e104. 10.7717/peerj-cs.104.

[40] Bankevich A, Nurk S, Antipov D, Gurevich AA, Dvorkin M, Kulikov AS, et al. SPAdes: A new genome assembly algorithm and its applications to single-cell sequencing. J Comput Biol 2012;19:455–77. 10.1089/cmb.2012.0021.

[41] Gurevich A, Saveliev V, Vyahhi N, Tesler G. QUAST: Quality assessment tool for genome assemblies. Bioinformatics 2013;29:1072–5. 10.1093/bioinformatics/btt086.

[42] Chklovski A, Parks DH, Woodcroft BJ, Tyson GW. CheckM2: a rapid, scalable and accurate tool for assessing microbial genome quality using machine learning. Nat Methods 2023 208 2023;20:1203–12. 10.1038/s41592-023-01940-w.

[43] Zankari E, Hasman H, Cosentino S, Vestergaard M, Rasmussen S, Lund O, et al. Identification of acquired antimicrobial resistance genes. J Antimicrob Chemother 2012;67:2640. 10.1093/JAC/DKS261.

[44] Moura A, Criscuolo A, Pouseele H, Maury MM, Leclercq A, Tarr C, et al. Whole genome-based population biology and epidemiological surveillance of Listeria monocytogenes. Nat Microbiol 2016;2:1–10. 10.1038/nmicrobiol.2016.185.

[45] Robertson J, Nash JHE. MOB-suite: software tools for clustering, reconstruction and typing of plasmids from draft assemblies. Microb Genomics 2018;4. 10.1099/mgen.0.000206.

[46] Liu B, Zheng D, Jin Q, Chen L, Yang J. VFDB 2019: a comparative pathogenomic platform with an interactive web interface. Nucleic Acids Res 2019;47:D687. 10.1093/NAR/GKY1080.

[47] Wishart DS, Han S, Saha S, Oler E, Peters H, Grant JR, et al. PHASTEST: Faster than PHASTER, better than PHAST. Nucleic Acids Res 2023;51:W443–50. 10.1093/nar/gkad382.

[48] Schwengers O, Jelonek L, Dieckmann MA, Beyvers S, Blom J, Goesmann A. Bakta: rapid and standardized annotation of bacterial genomes via alignment-free sequence identification. Microb Genomics 2021;7:685. 10.1099/MGEN.0.000685.

[49] ECDC and EFSA. Prolonged multi-country cluster of Listeria monocytogenes ST155 infections linked to ready-to-eat fish products. EFSA Support Publ 2023;20:8538E. 10.2903/SP.EFSA.2023.EN-8538.

[50] Croucher NJ, Page AJ, Connor TR, Delaney AJ, Keane JA, Bentley SD, et al. Rapid phylogenetic analysis of large samples of recombinant bacterial whole genome sequences using Gubbins. Nucleic Acids Res 2015;43:e15. 10.1093/nar/gku1196.

[51] Minh BQ, Schmidt HA, Chernomor O, Schrempf D, Woodhams MD, Von Haeseler A, et al. IQ-TREE 2: New Models and Efficient Methods for Phylogenetic Inference in the Genomic Era. Mol Biol Evol 2020;37:1530–4. 10.1093/MOLBEV/MSAA015.

[52] Letunic I, Bork P. Interactive Tree Of Life (iTOL) v5: an online tool for phylogenetic tree display and annotation. Nucleic Acids Res 2021;49. 10.1093/nar/gkab301.

[53] Moura A, Criscuolo A, Pouseele H, Maury MM, Leclercq A, Tarr C, et al. Whole genome-based population biology and epidemiological surveillance of Listeria monocytogenes. Nat Microbiol 2016;2:16185. 10.1038/NMICROBIOL.2016.185.

[54] Kastner R, Dussurget O, Archambaud C, Kernbauer E, Soulat D, Cossart P, et al. LipA, a Tyrosine and Lipid Phosphatase Involved in the Virulence of Listeria monocytogenes. Infect Immun 2011;79:2489. 10.1128/IAI.05073-11.

[55] Camejo A, Carvalho F, Reis O, Leitão E, Sousa S, Cabanes D. The arsenal of virulence factors deployed by Listeria monocytogenes to promote its cell infection cycle. Virulence 2011;2:379–94. 10.4161/VIRU.2.5.17703.

[56] Schmitz-Esser S, Gram L, Wagner M. Complete Genome Sequence of the Persistent Listeria monocytogenes Strain R479a. Genome Announc 2015;3:150–65. 10.1128/GENOMEA.00150-15.

[57] Parsons C, Lee S, Kathariou S. Dissemination and conservation of cadmium and arsenic resistance determinants in Listeria and other Gram-positive bacteria. Mol Microbiol 2020;113:560–9. 10.1111/MMI.14470.

[58] Corbett D, Schuler S, Glenn S, Andrew PW, Cavet JS, Roberts IS. The combined actions of the copper-responsive repressor CsoR and copper-metallochaperone CopZ modulate CopA-mediated copper efflux in the intracellular pathogen Listeria monocytogenes. Mol Microbiol 2011;81:457–72. 10.1111/J.1365-2958.2011.07705.X.

[59] Zapotoczna M, Riboldi GP, Moustafa AM, Dickson E, Narechania A, Morrissey JA, et al. Mobile-Genetic-Element-Encoded Hypertolerance to Copper Protects Staphylococcus aureus from Killing by Host Phagocytes. MBio 2018;9:e00550–18. 10.1128/MBIO.00550-18.

[60] Purohit R, Ross MO, Batelu S, Kusowski A, Stemmler TL, Hoffman BM, et al. Cu+-specific CopB transporter: Revising p1B-type ATPase classification. Proc Natl Acad Sci U S A 2018;115:2108–13. 10.1073/PNAS.1721783115/SUPPL_FILE/PNAS.1721783115.SAPP.PDF.

[61] Calhoun LN, Kwon YM. Structure, function and regulation of the DNA-binding protein Dps and its role in acid and oxidative stress resistance in Escherichia coli: a review. J Appl Microbiol 2011;110:375–86. 10.1111/J.1365-2672.2010.04890.X.

[62] Wagner J, Gruz P, Kim SR, Yamada M, Matsui K, Fuchs RPP, et al. The dinB gene encodes a novel E. coli DNA polymerase, DNA pol IV, involved in mutagenesis. Mol Cell 1999;4:281–6. 10.1016/S1097-2765(00)80376-7.

[63] Parsons C, Lee S, Jayeola V, Kathariou S. Novel Cadmium Resistance Determinant in Listeria monocytogenes. Appl Environ Microbiol 2017;83. 10.1128/AEM.02580-16.

[64] Hurley D, Luque-Sastre L, Parker CT, Huynh S, Eshwar AK, Nguyen S V., et al. Whole-Genome Sequencing-Based Characterization of 100 Listeria monocytogenes Isolates Collected from Food Processing Environments over a Four-Year Period. MSphere 2019;4. 10.1128/MSPHERE.00252-19.

[65] Moura A, Tourdjman M, Leclercq A, Hamelin E, Laurent E, Fredriksen N, et al. Real-Time Whole-Genome Sequencing for Surveillance of Listeria monocytogenes, France. Emerg Infect Dis 2017;23:1462. 10.3201/EID2309.170336.

[66] Zhang H, Chen W, Wang J, Xu B, Liu H, Dong Q, et al. 10-Year Molecular Surveillance of Listeria monocytogenes Using Whole-Genome Sequencing in Shanghai, China, 2009–2019. Front Microbiol 2020;11:551020. 10.3389/FMICB.2020.551020/BIBTEX.

[67] Surveillance Atlas of Infectious Diseases 2024. https://atlas.ecdc.europa.eu/public/index.aspx (accessed 26 June 2024).

[68] Committee for Antimicrobial Susceptibility Testing of the European Society of Clinical Microbiology E, Diseases I. Determination of minimum inhibitory concentrations (MICs) of antibacterial agents by broth dilution. Clin Microbiol Infect 2003;9:ix–xv. 10.1046/j.1469-0691.2003.00790.x.

[69] Koopmans MM, Engelen-Lee JY, Brouwer MC, Jaspers V, Man WK, Vall Seron M, et al. Characterization of a Listeria monocytogenes meningitis mouse model. J Neuroinflammation 2018;15:1–11. 10.1186/S12974-018-1293-3/FIGURES/4.

[70] Nüesch-Inderbinen M, Bloemberg G V., Müller A, Stevens MJA, Cernela N, Kollöffel B, et al. Listeriosis Caused by Persistence of Listeria monocytogenes Serotype 4b Sequence Type 6 in Cheese Production Environment. Emerg Infect Dis 2021;27:284. 10.3201/EID2701.203266.

[71] Maury MM, Tsai YH, Charlier C, Touchon M, Chenal-Francisque V, Leclercq A, et al. Uncovering Listeria monocytogenes hypervirulence by harnessing its biodiversity. Nat Genet 2016;48:308. 10.1038/NG.3501.

[72] Voronina OL, Kunda MS, Ryzhova NN, Aksenova EI, Kustova MA, Karpova TI, et al. Listeria monocytogenes ST37 Distribution in the Moscow Region and Properties of Clinical and Foodborne Isolates. Life 2023;13:2167. 10.3390/LIFE13112167/S1.

[73] Wang N, Yang L, Yuan Y, Wu C, He C. Clinical and Bacterial Characteristics of Bloodstream Infections Caused by Listeria monocytogenes in Western China. Can J Infect Dis Med Microbiol = J Can Des Mal Infect La Microbiol Médicale 2024;2024:7785327. 10.1155/2024/7785327.

[74] Wartha S, Bretschneider N, Dangel A, Hobmaier B, Hörmansdorfer S, Huber I, et al. Genetic Characterization of Listeria from Food of Non-Animal Origin Products and from Producing and Processing Companies in Bavaria, Germany. Foods 2023;12:1120. 10.3390/FOODS12061120/S1.

[75] Zhang Y, Dong S, Chen H, Chen J, Zhang J, Zhang Z, et al. Prevalence, Genotypic Characteristics and Antibiotic Resistance of Listeria monocytogenes From Retail Foods in Bulk in Zhejiang Province, China. Front Microbiol 2019;10:1710. 10.3389/FMICB.2019.01710.

[76] Pirone-Davies C, Chen Y, Pightling A, Ryan G, Wang Y, Yao K, et al. Genes significantly associated with lineage II food isolates of Listeria monocytogenes. BMC Genomics 2018;19. 10.1186/S12864-018-5074-2.

[77] Bolten S, Harrand AS, Skeens J, Wiedmann M. Nonsynonymous Mutations in fepR Are Associated with Adaptation of Listeria monocytogenes and Other Listeria spp. to Low Concentrations of Benzalkonium Chloride but Do Not Increase Survival of L. monocytogenes and Other Listeria spp. after Exposure to Benzalkonium Chloride Concentrations Recommended for Use in Food Processing Environments. Appl Environ Microbiol 2022;88. 10.1128/AEM.00486-22.

[78] Harvey J, Gilmour A. Characterization of Recurrent and Sporadic Listeria monocytogenes Isolates from Raw Milk and Nondairy Foods by Pulsed-Field Gel Electrophoresis, Monocin Typing, Plasmid Profiling, and Cadmium and Antibiotic Resistance Determination. Appl Environ Microbiol 2001;67:840. 10.1128/AEM.67.2.840-847.2001.

[79] Vu HTK, Benjakul S, Vongkamjan K. Characterization of Listeria prophages in lysogenic isolates from foods and food processing environments. PLoS One 2019;14. 10.1371/JOURNAL.PONE.0214641.

[80] Rostamian M, Kooti S, Mohammadi B, Salimi Y, Akya A. A systematic review and meta-analysis of Listeria monocytogenes isolated from human and non-human sources: the antibiotic susceptibility aspect. Diagn Microbiol Infect Dis 2022;102:115634. 10.1016/J.DIAGMICROBIO.2022.115634.

