## Supplemental Figures for "Genomic characterisation of *Listeria monocytogenes* isolated from normally sterile human body fluids in Lithuania from 2016 to 2021"

**Supplementary Figure S1**. Minimum Spanning Tree (MST) based on cgMLST scheme. Each circle represents genetically related isolates. The branch lengths between circles indicate the number of allele differences between connected circles. The size of each circle is proportional to the number of strains it contains. Colors correspond to the countries where the isolates were collected. Branches with more than 500 differences are truncated and depicted as dashed lines. Each cluster in **panel (A)** is labeled A - G in the figure, with expanded branches shown in **panel (B)** to illustrate all branches (allele distances of ≥1) between Lithuanian isolates and the nearest root node with close relatives.

**(A)**

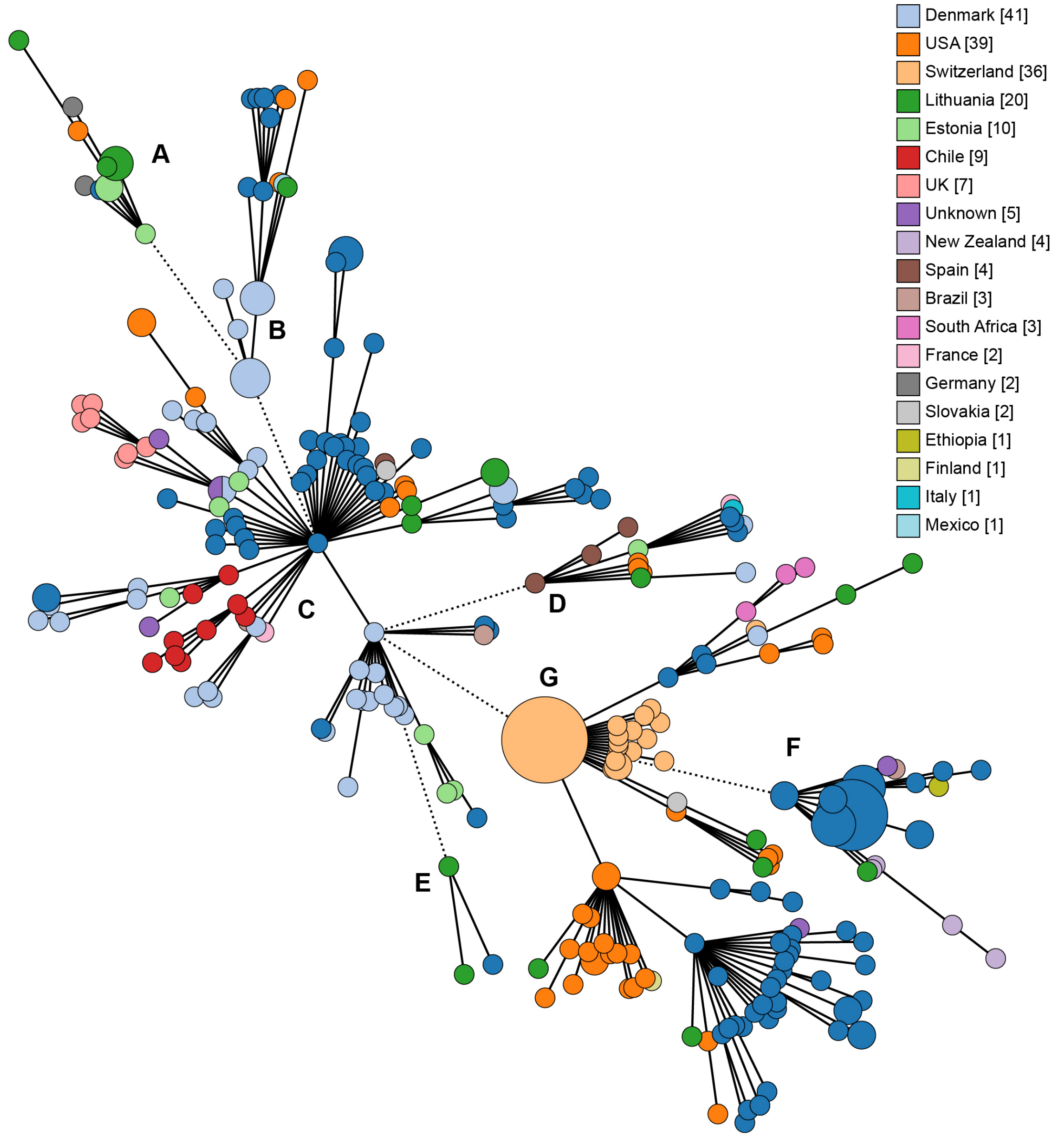

(B)

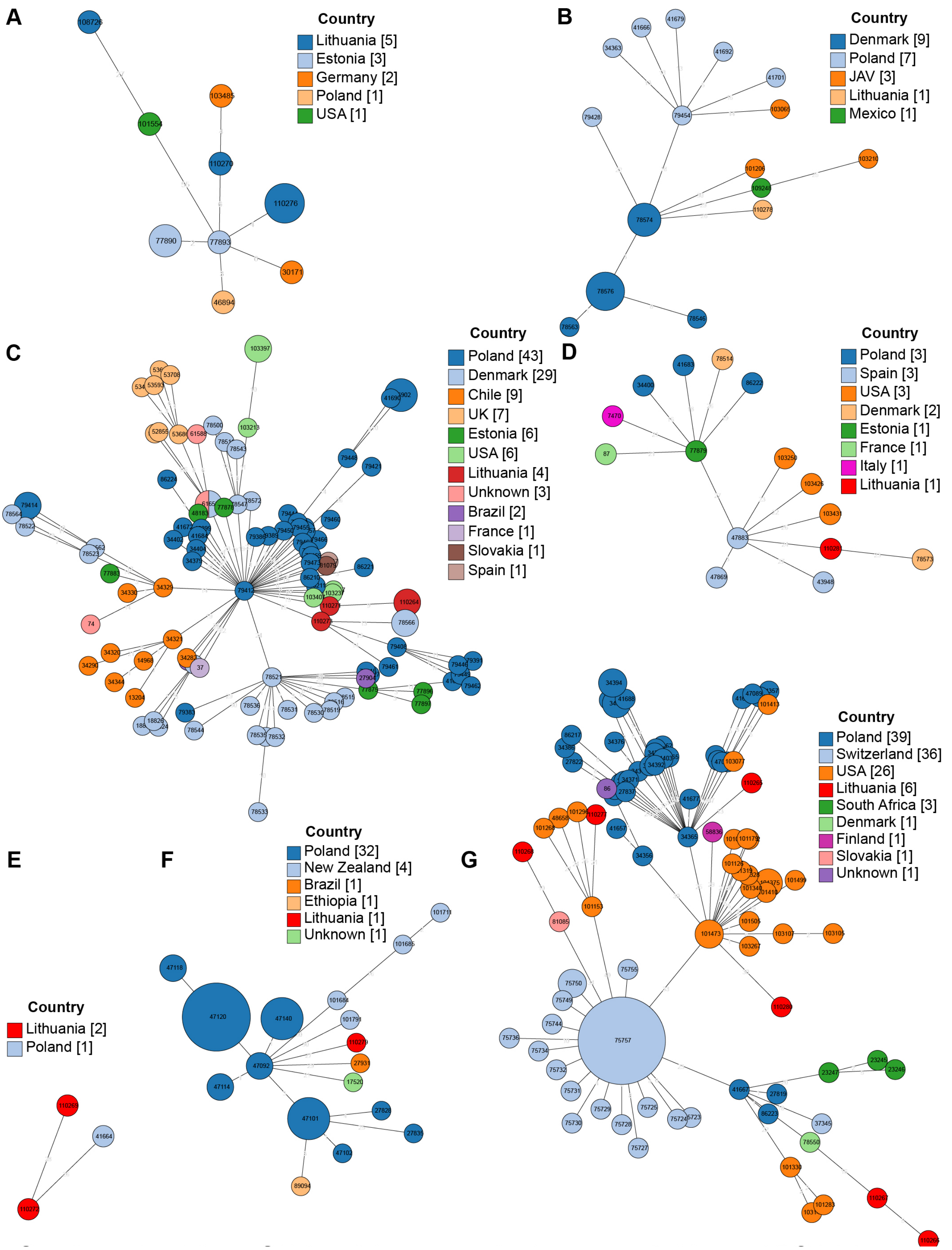
A
Country
Lithuania [5]
Estonia [3]
Germany [2]
Poland [1]
USA [1]
B
Country
Denmark [9]
Poland [7]
JAV [3]
Lithuania [1]
Mexico [1]
C
Country
Poland [43]
Denmark [29]
Chile [9]
UK [7]
Estonia [6]
USA [6]
Lithuania [4]
Unknown [3]
Brazil [2]
France [1]
Slovakia [1]
Spain [1]
D
Country
Poland [3]
Spain [3]
USA [3]
Denmark [2]
Estonia [1]
France [1]
Italy [1]
Lithuania [1]
E
Country
Lithuania [2]
Poland [1]
F
Country
Poland [32]
New Zealand [4]
Brazil [1]
Ethiopia [1]
Lithuania [1]
Unknown [1]
G
Country
Poland [39]
Switzerland [36]
USA [26]
Lithuania [6]
South Africa [3]
Denmark [1]
Finland [1]
Slovakia [1]
Unknown [1]

**Supplementary Figure S2.** Distribution of *L. monocytogenes* serotypes based on year of isolate collection **(A)** and age group **(B)**.

**(A)**

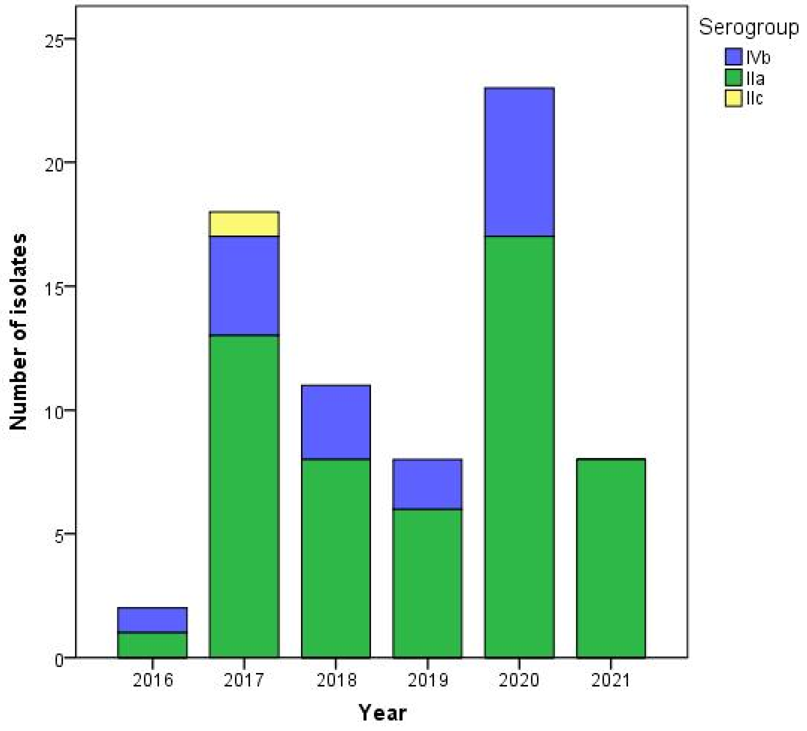


**(B)**

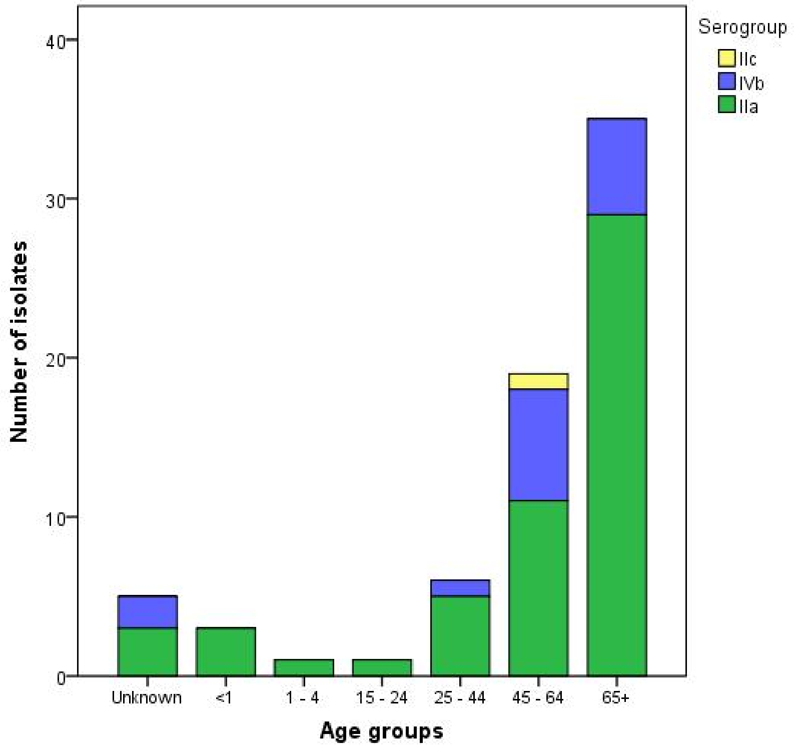
